# *Etmopterus* lantern sharks use coelenterazine as the substrate for their luciferin-luciferase bioluminescence system

**DOI:** 10.1101/2021.03.01.433353

**Authors:** Gaku Mizuno, Daichi Yano, José Paitio, Hiromitsu Endo, Yuichi Oba

## Abstract

The lantern shark genus *Etmopterus* is a group of deep-sea bioluminescent fishes. They emit blue light mainly from the ventral body surface, and the primary biological function is considered to be for camouflage by counterillumination. In this study, we detected both coelenterazine and coelenterazine-specific luciferase activity in the ventral photophore tissues. The results suggested that bioluminescence in lantern sharks is produced using coelenterazine as the substrate for the luciferin-luciferase reaction.

---

Bioluminescence in cartilaginous fish has been reported in species of the squaliform families Dalatiidae, Etmopteridae, and Somniosidae^1,2^. They have numerous tiny photophores mainly throughout the ventral body surface, and its biological function is considered to be counterillumination, a strategy for camouflage from predators by cloaking their silhouette and making them indistinguishable from environmental downwelling light^3,4^. Some of these species also possess photophores on the dorsal surface, fins and dorsal spines, and the light emission is suggested to have as an aposematic function or used for intraspecific communication^5,6^.

Coelenterazine, 6-(4-hydroxyphenyl)-2-[(4-hydroxyphenyl)methyl]-8-(phenylmethyl)-7H-imidazo[1,2-a]pyrazin-3-one, is a compound used as luciferin or chromophore of photoprotein in a wide range of marine bioluminescent organisms, including members of the phyla Radiozoa, Cercozoa, Porifera, Ctenophora, Cnidaria, Chaetognatha, Mollusca, Arthropoda, and Chordata^7,8^. Several coelenterazine-dependent luciferase or photoprotein genes have been isolated from ctenophores, *Aequorea* jelly, *Renilla* sea pansy, *Pholas* bivalves, *Sthenoteuthis* squid, *Oplophorus* shrimps and copepods, and some of these genes have been used as bio-tools in molecular biology^9,10^. Among bony fishes, more than 400 species of Stomiiformes and 250 species of Myctophiformes are considered to use coelenterazine-dependent bioluminescence systems^11–14^, but any of their luciferases have not yet been identified. Some luminous fish species in Pempheridae, Apogonidae, and Batrachoididae, which inhabit in shallow-water areas, are known to use cypridinid luciferin^9^; however, with the exception of *Parapriacanthus ransonneti*, which use cypridinid luciferase obtained from their ostracod prey^15^, their luciferase have not yet been determined.

In this study, we focused on the bioluminescence system employed by deep-sea sharks in the genus *Etmopterus* (Squaliformes, Etmopteridae). *Etmopterus* is one of the most diverse groups of sharks, with approximately 40 species currently recognized, and it is considered that they are probably all bioluminescent^3,16^. It has been reported that their bioluminescence systems are intrinsic (not symbiotic)^17,18^, but the reaction mechanism has not been elucidated. Based on our biochemical and chemical analyses, we concluded that etmopterid sharks use coelenterazine as the luciferin in their luciferin-luciferase bioluminescence system.

## Results

### Tissue distribution of luminescence activity in three *Etmopterus* species

Luminescence activities of various tissues in response to coelenterazine were examined in three species of *Etmopterus*: *E. molleri*, *E. pusillus*, *E. brachyurus*. The results showed that luminescence activity was predominant in tissues from the ventral skins for them (Fig. 1). Of note, Specimens 1 and 2 (Fig. 1A and 1B) are the same species morphologically of *E. molleri*, but they have small nucleotide differences between their *COI* sequences.

**Figure 1.**
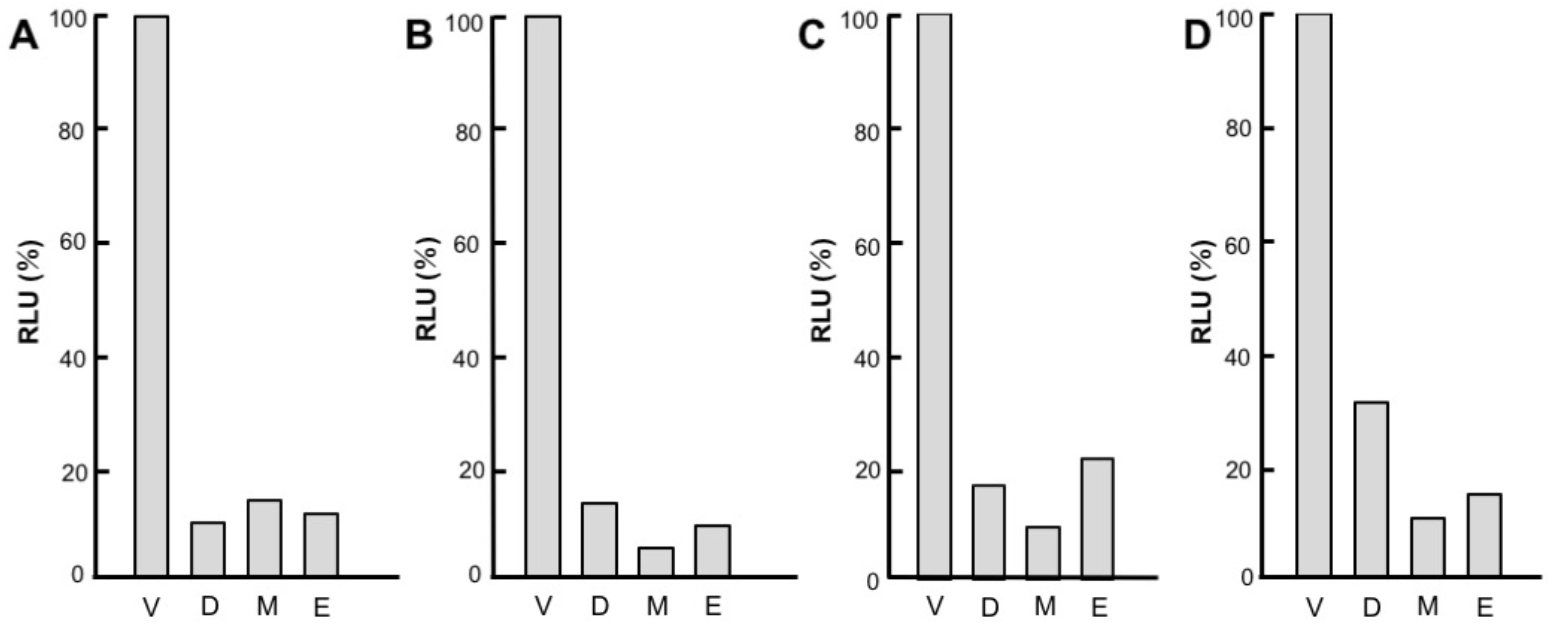
Tissue distributions of the luminescence activity with coelenterazine in *Etmopterus* spp. (A) *Etmopterus molleri* Specimen 1, (B) *E. molleri* Specimen 2 (C) *Etmopterus pusillus*, (D) *Etmopterus brachyurus*. V, ventral skin; D, dorsal skin; M, muscle; E, eye. The values are the mean of triplicate measurements and shown as percentage to values obtained for ventral skin.

### Detection of coelenterazine as active fraction

A crude methanol extract of ventral skin in *E. molleri* was separated using an octadecylsilane (ODS) column, and the luminescence activities of the fractions with the crude buffer extract were examined. The luminescence activity was detected as a single peak (Fig. 2A), and the retention time of the active fractions matches to that of authentic coelenterazine (Fig. 2B). Liquid chromatography-electrospray ionization-mass spectrometry/mass spectrometry (LC-ESI-MS/MS) analysis of the active fraction showed the molecular and fragment ion patterns (Fig. 2C) exactly agreement with those of coelenterazine (Fig. 2D)^19^. These results suggest that the crude methanol extract contained a compound that exhibited luciferin activity to intrinsic luciferase, and that the intrinsic luciferin is coelenterazine.

**Figure 2.**
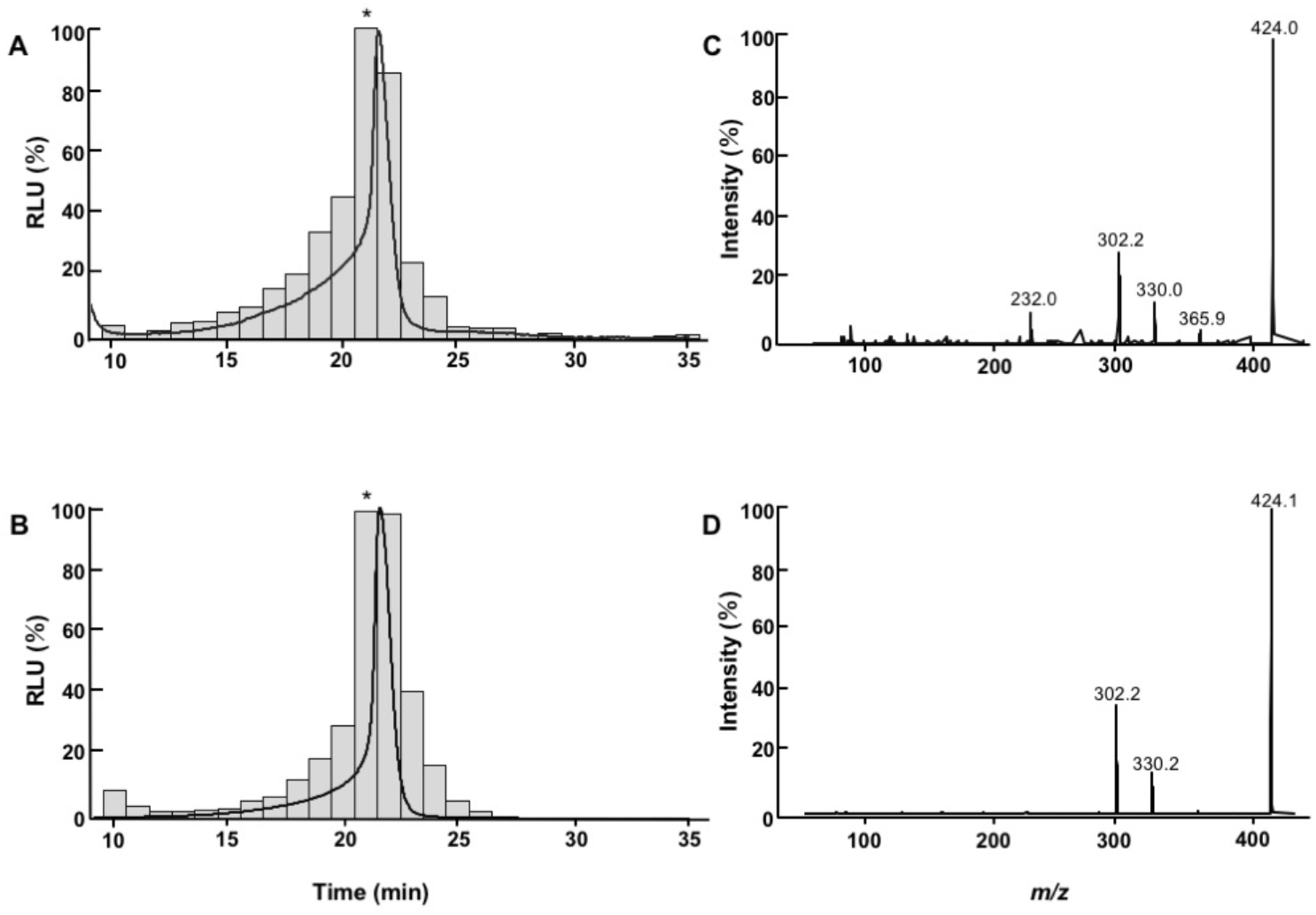
Luminescence activity and LC-ESI-MS/MS analyses of the HPLC fraction of the crude methanol extract from ventral skin in *E. molleri* (A, C) and authentic coelenterazine (B, D). (A, B) Lines represent fluorescence emission; bars represent the luminescence activity of the fraction. Asterisks represent the fraction used for the LC-ESI-MS/MS analyses. Panels C and D show the results of the product scan analysis for the fraction indicated by the asterisk of A and B, respectively.

### Luminescence active fraction is coelenterazine-specific enzyme

The crude buffer extract of the ventral skin in *E. molleri* was separated by a gel-filtration column, and the luminescence activities with crude methanol extract or authentic coelenterazine were examined. The results showed that the proteins in the crude buffer extract were separated effectively (Fig. 3A), and that the luminescence activity with crude methanol extract was detected as a single peak (Fig. 3B), the retention time for the activity peak matched that for the authentic coelenterazine (Fig. 3C). These results suggest that crude buffer extract contains a protein having luciferase activity to intrinsic luciferin, and the luciferase activity is coelenterazine specific.

**Figure 3.**
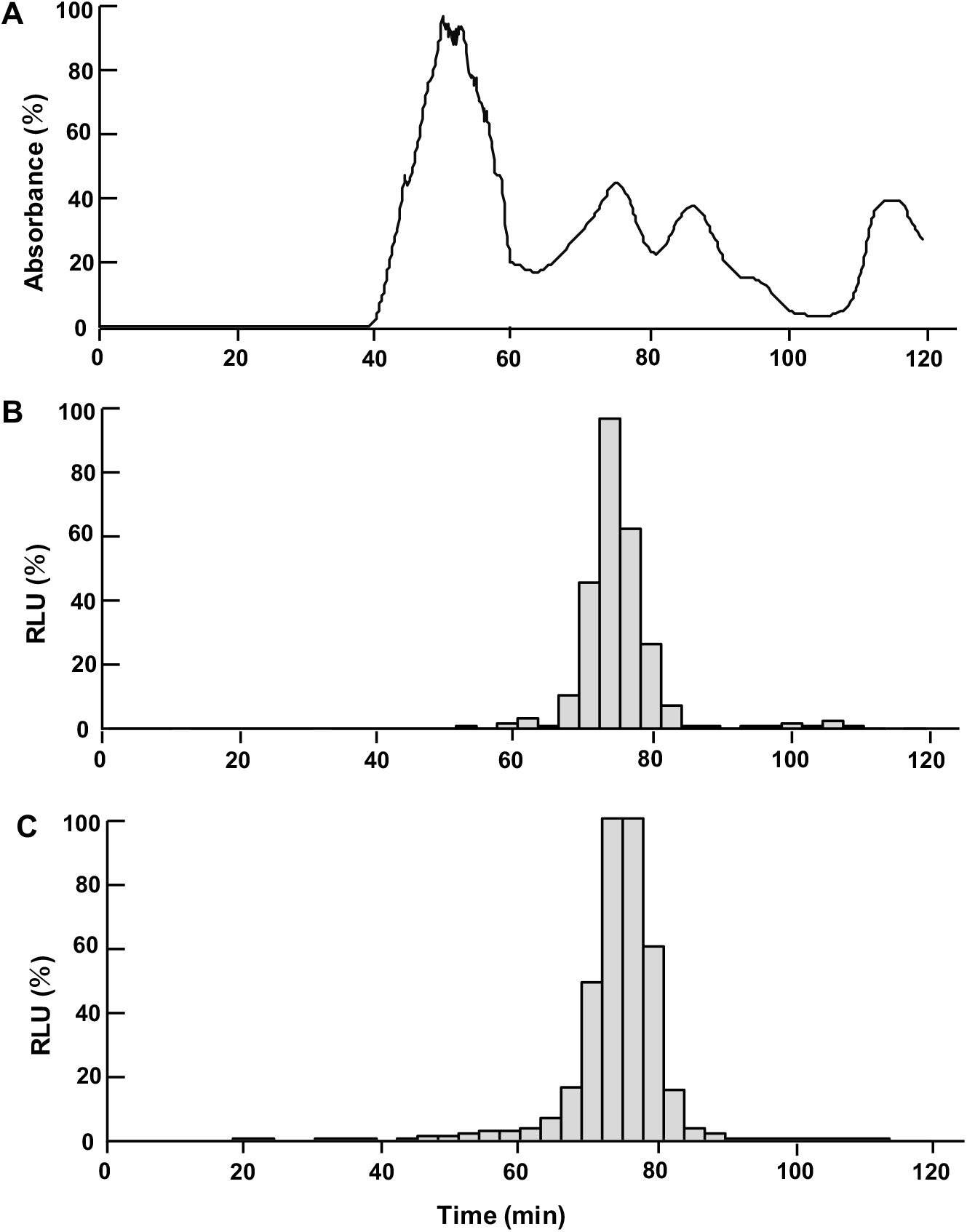
FPLC analyses of the crude buffer extract of ventral skin in *E molleri*. (A) UV detection at 280 nm. (B) Luminescence activity of the fraction with the crude methanol extract. (C) Luminescence activity of the fraction with authentic coelenterazine.

### Luminescence spectrum and pH dependence of the luciferase in *E. molleri*

The pH dependence of the luminescence activity was measured using the active fractions of the gel-filtration analysis of the crude buffer extract with authentic coelenterazine. The results showed that the optimum pH was around 7.0 (Fig. 4A). The luminescence spectrum at pH 7.0 showed an asymmetric single-peaked curve with a tail extending into the longer wavelength region, typical for those in the most bioluminescence organisms (Fig. 4B). The peak was at 460 nm, which corresponds to blue in human vision.

**Figure 4.**
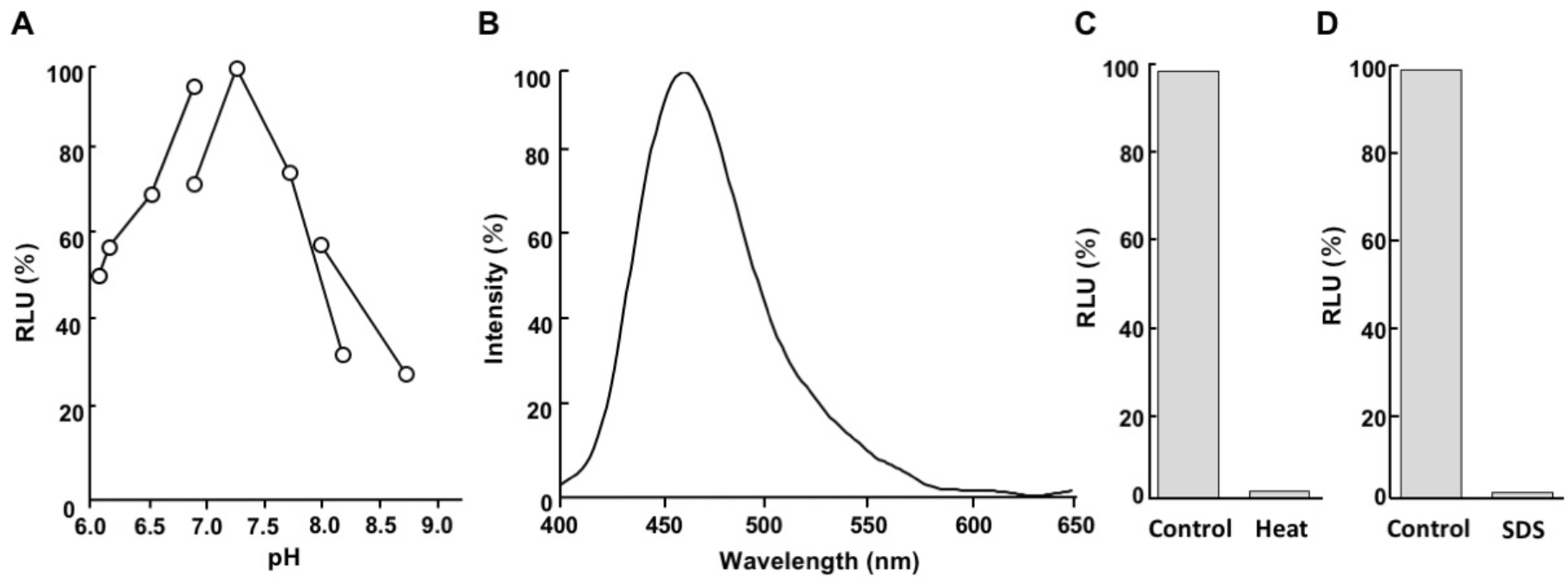
Enzymatic characteristics of the FPLC-purified luciferase with coelenterazine. (A) pH-dependent activity profiles. For pH adjustment, Bis-Tris-HCl, Tris-HCl, and glycine-NaOH buffers were used for pH 6-7, pH 7-8.5, and pH8.5-9.5, respectively. (B) Luminescence spectrum at pH 7.0. at 18 °C (C) Luminescence activity after the treatment at 18 °C for 15 min (Control) or 98 °C for 15 min (Heat). (D) Luminescence activity after the treatment without (Control) and with 2% SDS (SDS) at 18 °C for 5min.

### Heat and detergent susceptibility

The crude buffer extract from the ventral skin of *E. molleri* was treated by heat or sodium dodecyl sulfate (SDS) to test the enzyme susceptibility. The results showed that the luminescence activity was dramatically lost by the treatments (Fig. 4C and 4D). These results suggest that the luminescence activity in the buffer extract is attributed to protein and was denatured under non-physiological conditions like usual enzymes.

## Discussion

We extracted the luciferin activity by methanol and the luciferase activity by neutral buffer from lantern sharks, the genus *Etmopterus*. The luminescence activities of luciferase with coelenterazine were predominant in ventral skin compared to other tissues (dorsal skin, muscle, and eye) for three species of *Etmopterus*, *E. molleri*, *E. pusillus*, and *E. brachyurus*. These results are consistent with *in vivo* observations that the ventral light emission is much greater than dorsal light emission in *Etmopterus* species^6^. We then examined the luminescence mechanism in this genus using ventral tissues of *E. molleri*, as this species was collected most abundantly at Suruga Bay.

Identification of the luciferin molecule and luciferase protein is the most essential for understanding the bioluminescence mechanism of luminous organism being studied. In this regard, the Nobel laureate Osamu Shimomura stated “in studying bioluminescence reactions, it is crucially important to use the purified forms of the components necessary for light emission” as this avoids erroneous interpretations^9^. In the present study, therefore, we alternatively performed reciprocal analyses of the luciferin and luciferase using chromatographically separated fractions of luciferase and luciferin, respectively. The analyses confirmed that the luciferase activity in the buffer extract was coelenterazine-specific, and that the luciferin activity in the methanol extract was specific to coelenterazine-dependent luciferase. Mass spectrometry confirmed that the active fraction in the methanol extract is chemically identical to coelenterazine. The enzymatic activity was denatured both by heat and SDS treatments. The optimum pH of the observed luminescence activity was near physiological conditions, which is in contrast to the heterologous chemiluminescence activity observed for human albumin with coelenterazine at a highly alkaline pH^20^, suggesting that the luminescence activity in the shark skin extract is characteristic of a natural enzyme. The molecular size of the luciferase estimated by gel-filtration was 78-92 kDa. By further chromatographic separations, the luciferase protein can be purified.

The luminescence spectrum of the luciferase fraction with coelenterazine was peaked at 460 nm under neutral pH conditions. *In vivo* spontaneous luminescence spectra of the etmopterid shark species have been reported at wavelength of 476 nm in *Etmopterus splendidus*, 477 nm in *E. molleri*, and 486 nm in *E. spinax*^4^. The reason for the difference between our *in vitro* results and those obtained in that study is uncertain, but it may be due to genetic differences among the specimens or some effect of the colored pigment or reflectors in photophores.

Our findings contrast with those of a previous report suggesting no free-form of coelenterazine and no coelenterazine-dependent luciferase activities in the photophore tissues of *Etmopterus spinax*^21^. However, our findings showing that lantern sharks use coelenterazine for bioluminescence might be considered reasonable as lantern sharks inhabit the marine twilight zone and are higher up the food chain. It has been suggested that coelenterazine is biosynthesized in luminous copepods and comb jellies^22,23^, and that other luminous organisms generally obtained it through their diet, mainly by feeding on copepods or other organisms that contain coelenterazine^24^. During the dissection of the specimens, we found a variety of prey items in the stomach contents of *E. molleri*, including myctophid lanternfishes and unknown crustaceans. In analyses of *E. spinax* stomach contents, a variety of coelenterazine-dependent animals, such as *Oplophorus* and *Pasiphaea* shrimps, *Maurolicus* fish (Stomiiformes) and unidentified myctophid fishes, were also detected^21,25^. It is also known that some non-luminous fishes accumulated coelenterazine^26^. We consider that *Etmopterus* lantern sharks obtained coelenterazine for bioluminescence from fishes and crustaceans diets; they also obtained coelenterazine from their diets, such as copepods. In *E. spinax*, *in vivo* luminescence from ventral photophores was also detected in the late embryonic stage^27^, suggesting that coelenterazine is transferred vertically from the mother.

Bioluminescence in lantern sharks is considered to have originated in the Cretaceous Period^1^, which is coincidental with the origin of other coelenterazine-dependent fishes, Stomiiformes and Myctophiformes^28^. Identification of the luciferase gene and elucidation of the coelenterazine-transport system in lantern sharks, as well as in stomiiform and myctophiform fishes, might be helpful for understanding the evolution of deep-sea bioluminescence, how it appears to have occurred simultaneously in multiple fish lineages that inhabited deep-sea environment of the Cretaceous.

## Materials

### Shark samples

The specimens of *Etmopterus* were obtained from the by-caught of commercial bottom trawls at Suruga Bay, Japan in 2019 and 2020. All specimens were frozen at −20 °C after capture by T. A. Worldmedia (Shizuoka, Japan). The scientific name was identified by DNA analysis and morphological characteristics. The voucher specimens were deposited in the BSKU fish collection at Kochi University, Japan (*E. molleri*, BSKU 129578, 323 mm TL; *E. pusillus*, BSKU 129579, 201 mm TL; *E. brachyurus*, BSKU 129580, 280 mm TL). For DNA analysis, a partial region of the cytochrome *c* oxidase subunit I gene (*COI*) was amplified and sequenced. Briefly, total DNA was extracted from muscle tissue using Lysis Buffer for PCR (Takara, Shiga, Japan) and Proteinase K (Takara). Polymerase chain reaction (PCR) was performed using the primer set Fish-F1 and Fish-R2^29^ and Tks Gflex DNA polymerase (Takara, Japan). The PCR product was directly sequenced at Macrogen Inc. (Tokyo, Japan), and the sequence was deposited in the GenBank/ENA/DDBJ database (accession numbers, now depositing).

### Tissue distribution analysis

Each tissue sample (ventral skin, dorsal skin, dorsal body muscle, and eye ball) was dissected from a single defrosted specimen. Approximately 0.5 g of each tissue sample was homogenized in 500 μL of cold extraction buffer (20 mM Tris-HCl, 10 mM EDTA, pH7.4) using a plastic pestle, and centrifuged at 7,197 × *g* for 30 min. The extraction buffer (90 μL) containing 1.18 μM of synthetic coelenterazine (JNC Corporation, Tokyo, Japan) was injected into 30 μL aliquot of the supernatant, and the luminescence activity was measured for 2 min using a 96-well luminometer Centro LB960 (Berthold, Bad Wildbad, Germany). The accumulated value (RLU, relative light unit) was normalized by the protein concentration of the extract, which was measured using a Protein Assay Kit (Bio-Rad, Hercules, CA).

### Extraction and HPLC separation of luciferin

Dissected ventral skin samples of *E. molleri* were homogenized in cold methanol (mL/wet weight g) containing a 1:100 volume of 1M dithiothreitol using a homogenizer Ultra-Turrax T25 (IKA-Werke, Staufen, Germany), and centrifuged at 7,197 × *g* for 30 min. The supernatant was collected as the crude methanol extract. The crude methanol extract was desalted and concentrated using a MonoSpin C18 reversed-phase column (GL Science, Tokyo, Japan) and separated by high performance liquid chromatography (HPLC) (SEC System Prominence 501, Shimadzu, Kyoto, Japan) with a Cadenza CD-C18 column (2.0×75 mm, Imtakt, Kyoto, Japan). The fluorescence was detected at an excitation wavelength of 435 nm and an emission wavelength of 530 nm using a fluorescence detector RF-10AXL (Shimadzu). The mobile phase was an aqueous/methanol solution containing 0.1% formic acid, and the linear gradient of methanol was from 25% to 95% (2% per min). The flow rate was 0.2 mL/min. Fractions were collected at 1-min intervals.

### Extraction and FPLC separation of luciferase

Dissected ventral skin samples of *E. molleri* was homogenized in cold buffer (20 mM Tris-HCl, 10 mM EDTA, pH7.4) (1 mL/ wet weight g) using a homogenizer Ultra-Turrax T25, and centrifuged at 7,197 × *g* for 30 min. The supernatant was collected as crude buffer extract. The crude buffer extract was filtrated using a membrane filter Millex-SV 5.0 μm (Merck, Darmstadt, Germany) and separated by fast protein liquid chromatography (FPLC) using AKTA Prime Plus (Cytiva, Uppsala, Sweden). The column for gel filtration was a HiLoad 16/600 Superdex 200 prep grade (Cytiva) at a flow rate of 1.0 mL/min; mobile phase, 20 mM Bis-Tris-HCl, 150 mM NaCl, pH 7.4. Fractions were collected at 3-min intervals.

### Bioluminescence assay of HPLC fractions

A 30 μL aliquot of crude buffer extract in 265 μL of 20 mM Tris-HCl (pH7.2) was mixed with a 5 μL aliquot of each HPLC fraction, and the luminescence activity was measured for 5 min using a 96-well luminometer Centro LB960.

### Bioluminescence assay of FPLC fractions

A 100 μL of 1/300 diluted crude methanol extract or 1.18 μM coelenterazine in 20 mM Tris-HCl (pH7.2) was added into a 50 μL aliquot of each FPLC fraction, and the luminescence activity was measured for 5 min using a 96-well luminometer Centro LB960.

### Spectral measurement

*In vitro* bioluminescence spectrum was measured using the active fraction of the FPLC purification. The active fraction was concentrated using a 50 kDa cutoff filter Amicon Ultra-4 (Merck), and of 20 μL was mixed with 2 μL of 5.9 mM coelenterazine and 278 μL of 20 mM Bis-Tris-HCl (pH 7.0). The luminescence spectrum was measured using a fluorescence spectrophotometer FP-777W (Jasco, Tokyo, Japan) with the excitation light source turned off. The obtained raw spectrum was smoothed using a binomial method.

### Mass spectrometry

LC-ESI-MS/MS analysis was performed using the positive mode with nitrogen as the collision gas (collision energy, 30 V) using an API 4000 (AB SCIEX, Framingham, MA) connected to an LC800 HPLC system (GL Sciences) and a Cadenza CD-C18 column (Imtakt). The mobile phase was an aqueous/methanol solution containing 0.1% formic acid, and the linear gradient of methanol used was from 25% to 95% (2% per min). The flow rate was 0.2 mL/min. For the product ion scan analysis, an aliquot (5 μL) of HPLC fraction was applied and *m/z*= 424.0, corresponding to the calculated [M+H]^+^ mass value of coelenterazine, was monitored.

### Heat and SDS treatment

The crude buffer extract was heat treated using heat block MG-1200 (Eyela, Tokyo, Japan) set at 98 °C for 15 min, or mixed with SDS (final concentration, 2%) at room temperature (18 °C) for 5 min. A 90 μL of 1.18 μM authentic coelenterazine was injected into a 10 μL aliquot of heat or SDS treated extract, and the luminescence activity was measured for 1 min using a 96-well luminometer Centro LB960.

## Data availability

The authors declare that all data supporting the findings of this study are available within the article. DNA sequence data are deposited in GenBank/ENA/DDBJ.

## Author contributions

G.M., D.Y., and Y.O. conceived the idea of the study. Y.O. planned the study methodology. G.M., D.Y., J.P. and H.E. performed the experiments and species identifications. G.M. and D.Y. performed data analysis. G.M., D.Y., J.P., H.E. and Y.O. wrote the manuscript.

## Acknowledgements

We thank Prof. Kaname Tsutsumiuchi (Chubu University) for assistance with the HPLC and mass spectrometry measurements, and Assoc. Prof. Shiro Takei (Chubu University) for reviewing the manuscript and providing critical comments. The manuscript was proofread by a professional English editing service, Forte Co. This work was supported by JST CREST (JPMJCR16N1).

**Correspondence** and requests for materials should be addressed to Y.O.

